# EEG Signature of Idiopathic Hypersomnia: Insights from Sleep Microarchitecture and Hypnodensity Metrics

**DOI:** 10.1101/2025.05.26.656134

**Authors:** Arthur Le Coz, Raphael Vollhardt, Pierre Champetier, Ruben Herzog, Smaranda Leu-Semenescu, Pauline Dodet, Valérie Attali, Ana Gales, Delphine Oudiette, Isabelle Arnulf, Thomas Andrillon

## Abstract

**Background and Objectives:** Patients with idiopathic hypersomnia with long sleep time (IH) report daytime hypersomnolence despite prolonged sleep time and normal sleep macrostructure. As they often have non-restorative sleep, we investigated whether the structure of their sleep is abnormal.

**Methods:** In polysomnography recordings from 80 IH participants and 48 controls, we quantified hypnodensity metrics across the night (macro level), periodic and aperiodic spectral properties, infraslow fluctuations of sigma power within the night (meso level), slow waves, sleep spindles and their clustering (microstructure). Multivariate machine-learning models were used to classify IH vs. control sleep.

**Results:** Hypnodensity metrics were comparable between IH and controls, apart from more mixed wake/N1 sleep epochs in IH, and greater divergence between consecutive epochs of the same stage during NREM sleep in IH. Sigma power was increased in N2 sleep in IH and sleep spindles were more frequent and clustered. Slow wave density was higher in IH. Higher mean spindle cluster size correlated with higher Epworth Sleepiness Scale scores. Multivariate machine learning models incorporating these features achieved a balanced accuracy of 74% in distinguishing IH from controls.

**Discussion:** While spindles and slow waves are typically associated with good sleep quality, they are increased in IH patients. This could reflect greater need for sleep and increased difficulty waking up in IH, which is also characterized by more mixed wake/N1 stages.

## Introduction

Idiopathic hypersomnia is a rare, chronic disorder of hypersomnolence, characterized by excessive daytime sleepiness despite normal or prolonged sleep duration. The disorder is often associated with i) a feeling of unrefreshing sleep, ii) marked difficulty in waking up—called sleep drunkenness or severe morning inertia—and iii) long unrefreshing naps^1^. The diagnostic framework requires demonstration of excessive daytime sleepiness in the absence of sleep debt or other explanatory conditions, as evidenced by a mean sleep latency test (MSLT) latency ≤8 minutes (and fewer than two sleep onsets in REM) and/or an excessive sleep need, as evidenced by a total sleep time greater than 11 hours/24 hours of polysomnography monitoring. Two forms of idiopathic hypersomnia have been described based on different symptoms, measures, and clustering analysis^2^. Idiopathic hypersomnia with long sleep time (IH) is considered as an independent, rare clinical entity and is closely associated with sleep time>11h on polysomnography. In contrast, idiopathic hypersomnia without long sleep time is evidenced exclusively with short MSLT latency, seems to closely resemble narcolepsy type 2, and is more frequent (1.5%) in epidemiologic studies^3–6^. The cause of IH is unknown, but current hypotheses include a deficient arousal system (as in narcolepsy), a longer biological night^7,8^, the production of an endogenous hypnotic factor^9^, or an abnormal sleep structure.

The diagnosis of IH requires that sleep duration is normal or prolonged and that sleep is not excessively fragmented. In a meta-analysis of 10 studies, participants with IH had increased total sleep time and REM sleep percentage, decreased sleep onset latency and N3 percentage, and similar sleep efficiency and REM sleep latency compared to controls^10^. However, this meta-analysis mixed IH participants with and without long sleep time, and included studies with low sample size. In another recent large study, longer sleep time, shorter sleep onset latency and higher percentage of REM sleep were confirmed in the IH group, while sleep efficiency was higher and N3 percentage was unchanged compared to controls^11^. The arousal index was lower than in controls^11,12^. Overall, these studies suggest a longer and better consolidated sleep in IH, which contrasts with the complaint of excessive daytime sleepiness.

If the macrostructure of sleep is normal or even above normal, could an altered microstructure explain why sleep is not restorative in IH? Studies of microstructure in IH are scarce and yielded inconsistent results, potentially because they include heterogeneous IH groups with small sample sizes and usually compare IH and narcolepsy type 1 rather than healthy sleepers. Quantitative EEG analyses led to mixed results, with slow wave activity either shown to be lower in IH^13,14^, or on the contrary higher with a faster dissipation during the night^15^. Specifically, individuals with IH had a higher density of sleep spindles during N2 sleep than controls^16^ and individuals with narcolepsy^16,17^. The rate of cyclic alternating pattern (CAP, a marker of sleep instability) was lower in IH without long sleep time than in controls^13^. It is important to emphasize that these changes were typically subtle and tended toward better sleep in IH compared to narcolepsy type 1 and similar sleep microstructure between IH and controls.

Beyond the conventional sleep/wake classification and spectral EEG analysis, new fine-grained analyses of EEG signals have proven valuable in capturing subtle neural dynamics that better align with subjective assessments of sleep quality and quantity^18^. First, hypnodensities constitute a probabilistic approach of sleep scoring, in contrast with the standard discrete sleep hypnograms. By computing the probability of the 5 different sleep stages (Wake, N1, N2, N3 and REM sleep) using an automatic sleep scoring algorithm^19–21^, hypnodensities provide a good overview of hybrid sleep stages (i.e., epochs with a high probability of two or more sleep stages) and of the temporal variability of sleep stages^21^. Second, splitting the study of EEG dynamics between an aperiodic background activity and the (periodic) brain oscillations allow a more precise analysis of brain rhythms^22^. Third, recent studies have stressed the importance of infraslow modulations of sleep dynamics, with a time period of approximately 50 seconds during stable NREM sleep^23,24^. These infraslow fluctuations structure the temporal organization of sleep hallmarks such as sleep spindles^24^, and reflect modulations of noradrenergic activity during NREM sleep. Examining infraslow oscillations or the temporal clustering of sleep spindles could offer a window onto the neuromodulation of sleep and the period favorable for stage transition and arousals^25^. These aspects might be relevant for IH as patients experience enormous difficulty waking up from sleep and were never studied in this population. Therefore, we aimed to determine whether these new approaches to polysomnography analyses could reveal differences in IH and provide a more comprehensive understanding of its pathophysiology in a large, homogeneous group of participants with IH and controls. Our methodology included analyses of the macrostructure (hypnogram and hypnodensities), the mesostructure (power spectral density, periodic analysis, infra-slow activity), and the microstructure of sleep (slow waves, sleep spindles and their clustering). Lastly, we combined all EEG features extracted to classify IH vs. control sleep recordings using automated machine learning algorithms.

## Methods

### Participants

#### Participants with Idiopathic Hypersomnia

This is a retrospective study of clinical and polysomnography data collected in the Pitié-Salpêtrière Sleep Clinic, a National Reference Centre for Narcolepsies and Rare Hypersomnias, between July 2015 and February 2023 as part of the clinical diagnostic of IH. Criteria of inclusion for IH patients comprised: age > 18 years old, and diagnosis of IH as defined by international criteria: Patient had to experience daily irresistible urges to sleep or daytime sleep episodes for at least three months, without any presence of cataplexy. Polysomnography and the MSLT should not support a diagnosis of narcolepsy type 1 or 2^1^. At least one of the following conditions had to be met: the MSLT showed a mean sleep latency of less than or equal to 8 minutes, or the total sleep time over 24 hours was greater than or equal to 660 minutes, as measured by a 24-hour polysomnographic recording on night 2 and day 3. All patients had a standardized 48-hour sleep recording protocol, which included a first night of sleep with a 8-hour sleep opportunity window (night 1), followed by a MSLT with 5 nap opportunities (day 2), a second night and day of unrestricted sleep (*ad libitum*, night 2 and day 3 with two unlimited naps, stopped at 4 pm), which allowed measurement of unrestricted sleep time over a period of 18 hours^12,26^. Only the first night of sleep was analyzed in the present study. The diagnosis of IH was excluded if hypersomnolence or MSLT results were better explained by other sleep disorders, medical or psychiatric conditions, or the use of drugs or medications. Individuals were tested without sleep restriction to exclude a prolonged sleep duration due to chronic sleep deprivation, as confirmed by their sleep diary. Following international recommendations for MSLT, psychotropic medications, if present, were withdrawn for at least five half-lives for wake-enhancing drugs and stimulants, and at least three months for benzodiazepines, neuroleptics, antiepileptics, or antidepressants. This allowed to eliminate the effects of metabolites with long half-life (especially antidepressants) and potential rebound phenomena. For this study, only patients with IH with long sleep time (total sleep time >660 min/18h) were analyzed.

#### Control Group

We retrospectively analyzed polysomnographic data of the control group from two datasets. The first dataset consisted of healthy individuals who participated in a study of the effects of verbal violence on sleep (ethics committee-approved study, clinicaltrial.gov reference: NCT03074578). These controls spent two nights in the sleep clinic, but we analyzed only the first (baseline) night, which was obtained under the same conditions as night 1 of the 48-hour protocol of the IH patients. The second data set consisted of young adults who were referred to the sleep clinic for a systematic screening for sleep apnea syndrome during the preoperative evaluation of facial dysmorphia. As part of their evaluation, they underwent one night of polysomnography under the same conditions as night 1 of the 48-hour protocol. Participants with a sleep time lower than 360 min, a sleep efficiency lower than 75%, a periodic leg movement index greater than 15 and an apnea-hypopnea index greater than 10 were excluded. All participants underwent a medical interview with a sleep specialist to determine the absence of any medical, sleep, or psychological complaints. Only individuals not taking any psychotropic medications or drugs were selected. Other exclusion criteria included the presence of another sleep disorder (e.g., parasomnias, all polysomnograms include a video-audio recording that is analyzed by the physician, or restless legs syndrome), comorbid neurological or psychiatric disorders, moderate to severe depressive symptoms, sleep debt prior to sleep testing as assessed by sleep diary, interview, or current or past night or shift work.

#### Ethics

Patients and preoperative control participants did not object to the anonymous reuse of their clinical and polysomnographic data for future research, as documented in their medical records, collected as part of their routine care and stored in a database declared to the National Commission for Data Protection and Liberties, in accordance with French law. Healthy controls were recruited from individuals who attended the sleep clinic as part of an ethics committee-approved research protocol and provided written informed consent, including permission for the anonymized reuse of their data.

#### Clinical measures

An experienced neurologist collected detailed information from the medical records of both patients and controls. These records included data obtained during a face-to-face interview with a sleep specialist, results from clinical scales, and a systematic self-report questionnaire completed upon entry to the sleep laboratory. The collected information covered: age, sex, body mass index, usual sleep and wake times during weekdays and weekends, presence of daily naps, occurrences of sleep drunkenness, sleep attacks, hypnagogic hallucinations, and sleep paralysis. Additionally, data on Epworth Sleepiness Scale scores at the time of the polysomnography, ongoing treatments, recreational drug use, depressive symptoms, and comorbid sleep, medical, neurological, and mental disorders were recorded.

#### Polysomnography

The polysomnography was conducted following the American Academy of Sleep Medicine guidelines^27^. The EEG montage included three electrodes at frontal (F3), central (C3), and occipital (O1) sites, contralateral mastoid (A2), and a ground channel on the forehead. Additional channels included left and right electrooculography and surface electromyography from the chin muscle and left and right tibialis anterior muscles. These channels were recorded at a minimum sampling frequency of 256Hz using a Grael amplifier (Compumedics Ltd, Australia). Cardiorespiratory measures included nasal pressure, nasal-oral thermistor airflow, tracheal sound sensor, thoracic and abdominal respiratory inductance plethysmography, position sensor, pulse oximetry, and electrocardiographic DII derivation. All recordings featured infrared video monitoring and ambient room sound recording. Sleep stages, micro-arousals, respiratory and motor events were scored on 30s epochs by certified sleep specialists according to the manual version 2.4^28^, using Profusion Sleep 4 software (*Compumedics Ltd, Australia*).

### Data Analysis

#### Preprocessing

We analyzed the EEG (F3, C3, and O1), electromyography (chin derivation), and electro-oculography channels. All channels were resampled to a common sampling rate of 256 Hz. To minimize the impact of electrical noise, data were notch-filtered at 50 Hz and 100 Hz using a zero-phase notch filter. A zero-phase finite impulse response band-pass filter was then applied to retain frequencies between 0.5 and 40 Hz. The EEG signals were re-referenced to the opposite mastoid (A2) electrode, providing a common reference point across recordings.

#### Automated Sleep Staging Algorithm and Extraction of Hypnodensity

Traditional sleep scoring by human experts is the gold-standard method for the identification of the different sleep stages. However, scoring is discrete and can vary between individual scorers^29,30^. The YASA package^20^ was used here for the automated sleep staging but other existing algorithms could be used (see^31^ for a benchmark). For each 30s-epoch, the YASA algorithm computes a probability of each stage (wake, N1, N2, N3 and REM sleep), providing both hypnodensities (graded evaluation of sleep stages^19,20^) and a standard hypnogram (by selecting the stage with the highest probability for each epoch). To evaluate the agreement between expert sleep scoring and YASA’s automated scoring, we computed Cohen’s kappa using the scikit-learn package^32^. We also computed confusion matrices to compare the expert and automated sleep scoring.

#### Intrusions and Variability

To characterize sleep dynamics, we employed two information theory measures: entropy (H) and Kullback-Leibler divergence (*D_KL_*)^33^. These metrics were computed at the epoch level based on the distribution of hypnodensities (probability of Wake, N1, N2, N3 and REM) to quantify the scoring ambiguity (which can be due to stage mixing^34^) and epoch-to-epoch variability, respectively. To maximize the interpretability of hypnodensities, and the derived metrics, we analyzed in conjunction with human expert scoring.

Entropy (H) measures the uncertainty associated with a probability distribution:

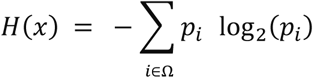

Where x represents a given epoch, Ω represents the five possible stages (wake, N1, N2, N3, REM sleep), and *p_i_* is the probability of each stage. Higher entropy indicates greater stage ambiguity, potentially reflecting intrusions from other stages.

To measure variability between consecutive epochs, we used *D_KL_*: For two probability distributions P(x) and Q(x) (i.e., the hypnodensity of two consecutive epochs) the *D_KL_* follows:

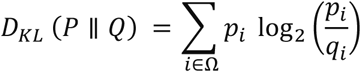

Where *p_i_*and *q_i_* is the probability of the stage i for the P and Q hypnodensity, respectively.

A *D_KL_* of zero indicates minimal variability (identical distributions), while higher values suggest greater differences between epochs in the distribution of hypnodensities. To examine the epoch-to-epoch variability of sleep stages above and beyond stage transitions, *D_KL_*was computed only on consecutive epochs of the same sleep stage according to the automated sleep scoring.

#### Spectral Analysis

Spectral analyses were performed on raw EEG data using the MNE-Python library^35^. We excluded wake periods before and after the night sleep (“lights out” period), as well as epochs containing micro-arousals. Power Spectral Density (PSD) was computed for channels F3, C3, and O1 on each epoch using the Welch method with a Hamming window (0.5-40 Hz frequency range, 1024-point Fast Fourier Transform (FFT) with 50% overlap). The PSD of the EEG signal can be decomposed into its periodic and aperiodic components^22^. The periodic component is thought to reflect brain oscillations (e.g., alpha oscillations, sleep spindles), and the aperiodic component the non-oscillatory background activity. We used the FOOOF package^22^ to model these periodic and aperiodic components with fixed aperiodic mode and peak width limits set to (0.5, 4) Hz. Before fitting FOOOF, we applied locally weighted scatterplot smoothing (LOWESS, span=0.075) to the PSD of each epoch to improve the robustness of the fit.

The aperiodic component follows a power law:

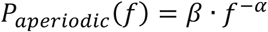

Where P_aperiodic_(f) is the aperiodic power for a given frequency f, α is the exponent of the power law, and β the offset. Subtracting the fitted aperiodic activity to the original PSD allowed to estimate the periodic power:

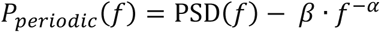

Absolute and relative periodic band power were computed for five frequency bands: delta (1–4 Hz), theta (4–8 Hz), alpha (8–12 Hz), sigma (12–16 Hz), and beta (16–30 Hz). For each 30s epoch, absolute band power was estimated by integrating the power spectral density within each frequency range in the periodic power spectrum. Relative band power was then computed by normalizing each band’s absolute power by the total power across all bands. All band power values were log-transformed to approximate a normal distribution.

#### Sleep Spindle and Slow Wave Automatic Detection

Sleep spindles and slow waves were automatically detected using custom Python scripts incorporating the MNE^35^ and YASA^20^ packages. For both spindles and slow waves, event density was calculated as the number of detected events within each sleep stage, divided by the total duration of that stage. To exclude wake periods unrepresentative of resting states, the analysis excluded wake epochs before and after sleep (lights-out and final awakening).

Slow waves were identified in all sleep stages, following a two-step preprocessing pipeline. First, EEG signals were down-sampled to 100 Hz and filtered using a zero-phase finite impulse response band-pass filter (0.5–8 Hz) with automatically determined transition bandwidths, ensuring minimal edge artifacts while preserving waveform integrity. The filtered signal was then analyzed to identify negative and positive zero-crossings, which were used to segment potential slow waves. For each candidate wave, we computed the peak-to-peak amplitude, that is the amplitude difference between the most positive and most negative peak between the start and end of the wave (consecutive zero-crossings). We then selected high-amplitude slow waves based on this peak-to-peak amplitude, which had to exceed a data-driven threshold, estimated for each channel and each participant. To compute this threshold, we fitted an exponentially modified Gaussian distribution (Ex-Gaussian) distribution to the distribution of peak-to-peak amplitudes for this channel, session and participant (scipy.stats^36^). This Ex-Gaussian fit provided 3 parameters: the mean (μ) and standard deviation (σ) of the Gaussian component and the Exponential tail (τ). To capture the tail of the Ex-Gaussian distribution, we defined the peak-to-peak threshold as twice the amplitude of the peak of the Ex-Gaussian. In N2 sleep and REM sleep, waves with PTP < 150µV were retained, while in N3 sleep, a higher threshold of 250µV was used to accommodate the larger amplitude of slow waves in N3 sleep. For each slow wave, we further extracted the following parameters: (1) Maximal negative slope (descending phase), representing the voltage decline (in µV/s) between the first negative zero-crossing and the negative peak crossing divided by the duration between the negative zero-crossing and the negative peak (in seconds); and (2) Maximal positive slope (ascending phase), reflecting the voltage increase between the negative peak and the subsequent positive zero-crossing divided by the duration between the peak and the positive zero-crossing (in seconds). Only waves with a descending and ascending slope between 0.25 and 2 µV/s were retained, ensuring the selection of physiologically meaningful slow waves. Slow waves were detected in each 30s epoch for EEG channels F3, C3, and O1, and stored for further analysis.

Sleep spindles were identified in N2 sleep and N3 sleep using the YASA spindle detection algorithm^20^, which applies both time-domain and frequency-domain criteria. EEG signals were first filtered between 12 and 15 Hz using a zero-phase FIR band-pass filter with a 1.5 Hz transition bandwidth on each side, ensuring that the -6 dB cutoff points were at 11.25 Hz and 15.75 Hz. To enhance specificity, spindles were detected within a broadband range of 1–30 Hz, which allowed for the computation of relative spectral power. Detected spindles were required to meet the three following morphological and spectral thresholds. First, duration constraint: events had to last between 0.5 and 2.5 seconds. Second, minimum inter-spindle interval: spindles occurring less than 500ms apart were merged into a single event. Third, two signal-based thresholds: (i) Root Mean Square threshold, where spindles had to exceed 1.5 standard deviations above the mean Root Mean Square of the 12–15 Hz filtered signal; (ii) Moving correlation threshold, requiring a correlation of ≥ 0.65 between the broadband and spindle-band filtered signals. Spindles were further characterized based on their peak-to-peak amplitude (µV), Root Mean Square power (µV), absolute power (log-transformed µV²), relative power (proportion of power in the spindle band relative to broadband), frequency (Hz), number of oscillations (count of positive peaks per event), and symmetry (location of the most prominent peak within the spindle, normalized between 0 = start and 1 = end).

Additionally, spindle clusters were analyzed using custom Python scripts. Clusters were defined as contiguous spindles separated by ≤6 seconds. Cluster size was defined as the number of spindles in each bout. Inter-cluster interval was computed as the time difference between the last spindle of a cluster and the first spindle of the subsequent bout, with temporal distances ≥300 s discarded. For each subject and channel, total spindle counts and the percentage of clusters for sizes 1 to ≥5 were computed.

#### Infraslow Rhythms

We adapted our analysis of fast spindle band power fluctuations from the literature^37,38^. Briefly, for each participant, we selected consecutive 120-s periods of artifact-free N2 and N3 sleep. Time–frequency power was computed in 100-ms bins with 0.2 Hz steps using a continuous wavelet transform with Morlet wavelets (four-cycle length). The resulting power time courses were smoothed with a symmetric 4-s moving average and normalized to the average absolute power in the sigma band (12 to 16 Hz) across all NREM sleep epochs. For each bout, power spectra in the 0.5–24 Hz range were computed every 0.1 s in steps of 0.2 Hz using a four-cycle Morlet wavelet transform, and the average power in conventional frequency bands (including the sigma band, 11–16 Hz) was determined. These power time courses were then smoothed using a 4-s symmetric moving average to highlight infraslow periodicities and normalized using the average absolute power computed over all N2 and N3 sleep epochs during the night. Finally, the spectral profiles of these power time courses were obtained via a continuous wavelet analysis in the 0.001–0.12 Hz range (with 0.001 Hz resolution) and averaged across bouts weighted by their duration.

#### Machine Learning Algorithm

We used machine learning models both for classifying the polysomnography of IH participants vs controls, and to regress sleep parameters based on EEG-derived features. To enhance interpretability and reduce noise in our high-dimensional EEG dataset, we applied a feature selection procedure based on the Minimum Redundancy Maximum Relevance algorithm^39^. For both classification and regression tasks, MRMR ranked the 26 EEG features by evaluating their relevance—using a random forest performance metric—and penalizing redundancy, which was quantified as the mean correlation with all other features (see Supplemental Table 1 for the list of features used). Features were then progressively incorporated into the prediction models according to their Minimum Redundancy Maximum Relevance ranking. For each subset, cross-validated (5-fold validation scheme) performance was computed (balanced accuracy and confusion matrices for classification and root-mean-square error for regression). The optimal feature subset was determined as the smallest set beyond which additional features did not yield statistically significant performance improvements (assessed via a Wilcoxon rank-sum test). For classification, Minimum Redundancy Maximum Relevance followed by repeated cross-validated modelling on both the full sample (n=128; 48 CT, 80 IH) and a balanced subsample obtained by bootstrapping the IH group 30 times. All classification and regression analyses were conducted using the XGBoost algorithm^40^, which builds a series of decision trees in a boosting framework, offering strong performance and robustness in high-dimensional datasets. For diagnostic group classification, we employed a softprob objective function with logloss as the evaluation metric, a learning rate of 0.1, a maximum tree depth of 3, 80% subsampling for both subjects and features, and a gamma regularization parameter of 0.1. In regression tasks aimed at predicting continuous sleep quality metrics (e.g., sleep efficiency, total sleep time, wake after sleep onset), the objective function was set to minimize Root Mean Square Error while retaining the same parameter settings. Model performance was rigorously evaluated using a repeated 5-fold cross-validation procedure (typically 60 training-testing iterations). In each iteration, data were partitioned into five balanced and stratified folds, with the model trained on four folds and validated on the remaining fold. This systematic procedure ensured that performance metrics—including balanced accuracy for classification and Root Mean Square Error for regression— were robust and generalizable across the dataset.

#### Statistics

We employed linear mixed-effects models to analyze sleep-related metrics, using the *statistical models* package in Python^41^. These models were designed to account for individual variability by including random effects based on subject identifiers. The linear mixed effects statistics were used on several EEG features including slow waves density, sleep spindle density, hypnodensities features (entropy, *D_KL_*, distribution), and scoring confusion matrices. These features were computed at the level of each epoch and averaged across epochs of a given sleep stage or directly computed for a given sleep stage. For each metric, the models included fixed effects for age and gender and interactions between sleep stage, group, and channel, where applicable. To ensure the robustness of our findings, we applied the false discovery rate method to correct for multiple comparisons^42^. However, for the power analysis, where p-values were computed for each frequency bin (0.25 Hz bin), we employed a cluster-based multiple comparison correction, which better accounts for the inherent spectral structure by identifying significant clusters of adjacent frequencies^43^. Additionally, demographic characteristics were analyzed using non-parametric tests, including the Mann-Whitney U test and Chi-squared test, to compare groups on variables such as age, body mass index, and scores at the Epworth sleepiness scale.

## Results

### Clinical and Sleep Characteristics of the Groups

Of the 384 patients diagnosed with idiopathic hypersomnia during the study period, 289 were excluded because they had an idiopathic hypersomnia without long sleep time (N = 14), were taking a psychotropic treatment (N = 123) or an illicit substance (N = 4), had moderate to severe depressive symptoms (N = 95), had a comorbid sleep, neurological, medical or psychiatric disorder (N = 141), slept less than 360 minutes on night 1 and/or had a sleep efficiency of less than 75% (N = 38), and/or their polysomnography was not accessible (N = 1). Thus, 80 participants with IH (with long sleep time) and 48 control participants were included in the analysis (Table 1). The total sleep time was longer than 11h/18h of recording in all 80 patients with IH including a total sleep time of 626.4 ± 102.9 min during night 2 and 715.5 ± 78.5 min during 18h recording. The mean MSLT latency was 11 ± 4.5 min in the IH group, and 21/80 (26.3%) participants had an MSLT latency ≤ 8 min. The prevalence of females was higher in the IH group than in the control group, but age and body mass index did not differ between groups. As expected from the characteristics of IH, usual weekday and weekend sleep times were longer in the IH than in the control group, and IH participants were more likely than controls to report sleep drunkenness, sleep attacks, daily naps, hypnagogic hallucinations and sleep paralysis. The Epworth sleepiness scale score was higher in the IH group than in the control group. During the first night of polysomnography (which was stopped at 6:30 AM in all groups), IH had higher sleep efficiency and N3 percentage, but lower wake after sleep onset and REM percentage compared to controls. The other measures, including sleep latencies, total sleep time, N1 and N2 percentage, as well as arousal index, periodic leg movement index and apnea-hypopnea index did not differ between groups.

### Automated Sleep Staging and Hypnodensities

The hypnogram was computed with the YASA algorithm and compared to the expert scoring. There was a good overall agreement (**K** = 0.693 ± 0.009 – Supplemental Figure 1) even if the YASA algorithm had not been trained on IH data^20^. The agreement between expert and YASA scoring for REM sleep was significantly higher in the IH group than in the control group (Figure 1b, ß = 0.054, p = 0.006), indicating more accurate automated detection of REM in IH patients. The entropy values of hypnodensities did not differ between groups. The epoch-to-epoch variability in these distributions, assessed by *D_KL_* showed a higher variability in the IH group (larger *D_KL_*) than in control group during N1 sleep (β = 0.041, p = 0.043), N2 sleep (β = 0.043, p = 0.042), and N3 sleep (β = 0.046, p = 0.042) epochs (Figure 1d). The hypnodensity-derived probability distributions of automated vs. traditional expert sleep scoring had similar profiles in IH and control groups (Supplemental Figure 1b). This suggests that hybrid sleep states—where features of multiple sleep stages overlap—were as prevalent in participants with IH as in controls. However, using the YASA scoring as a reference (Supplemental Materials, Figure 1a), the participants with IH have a lower probability of scoring wake during wake (0.78 < 0.82, ß = -0.0375, p = 0.0004) and a higher probability to score N1 sleep during wake (0.15 > 0.12, ß = 0.026, p = 0.00995) than controls. During N3 sleep, there was a higher probability for YASA to score N3 in participants with IH than in controls (0.85 > 0.84, ß = 0.016, p = 0.037).

**Figure 1:**
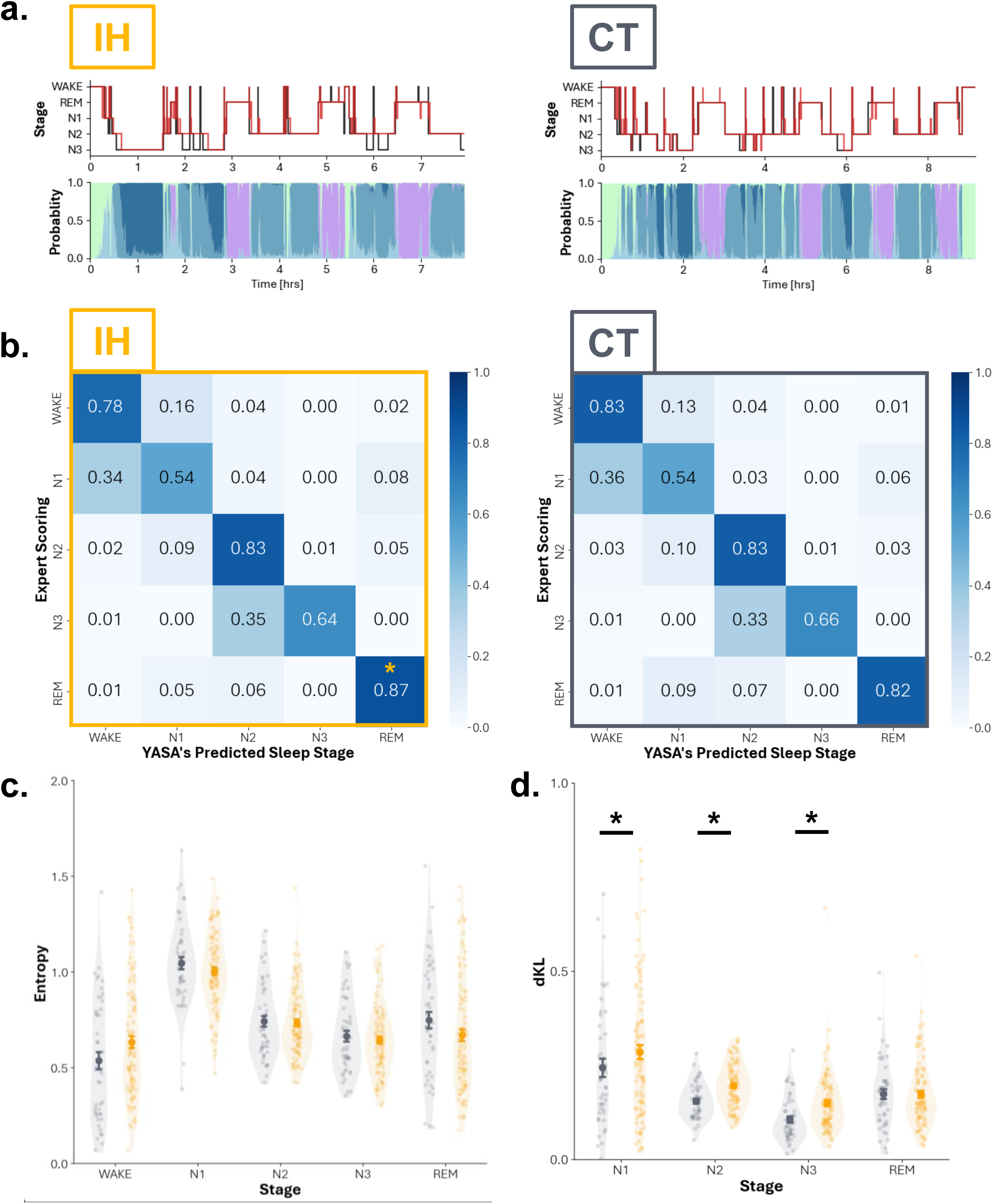
Participants with idiopathic hypersomnia do not have hybrid sleep states. **a** On the top row, hypnogram from experts scoring (black), and YASA scoring (red). On the bottom row, hypnodensities obtained with YASA in participants with IH (left panel) and controls (right panel). **b** Confusion matrices (experts vs YASA scoring). **c** Entropy (H) and epoch-to-epoch variability (D_KL_) extracted on YASA’s hypnodensity in the different sleep stages, participants with IH (yellow) and controls (grey). Statistics: linear mixed models with age and gender as covariate, correction for multiple comparisons (false discovery rate), p < 0.05: *, p < 0.01: **, p < 0.001: ***.

### Periodic Sigma Power

When comparing the EEG dynamics between groups, including the periodic and aperiodic power across sleep stages, there was an increase in the periodic power within the sigma band over the occipital (cluster: [12.5-17]Hz, p_CLUSTER_ < 0.001) and central electrode (cluster: [12.5-15.75]Hz, p_CLUSTER_ < 0.001) in N2 sleep in the IH group compared to the control group (Figure 2b). A similar increase of the frequency comprised between 12 and 18 Hz (which include the sigma band) was found in N1 sleep in all electrodes in the IH group compared to the control group (supplementary, Figure 3b p < 0.001). Other metrics of the periodic and aperiodic PSD did not differ between groups.

**Figure 2:**
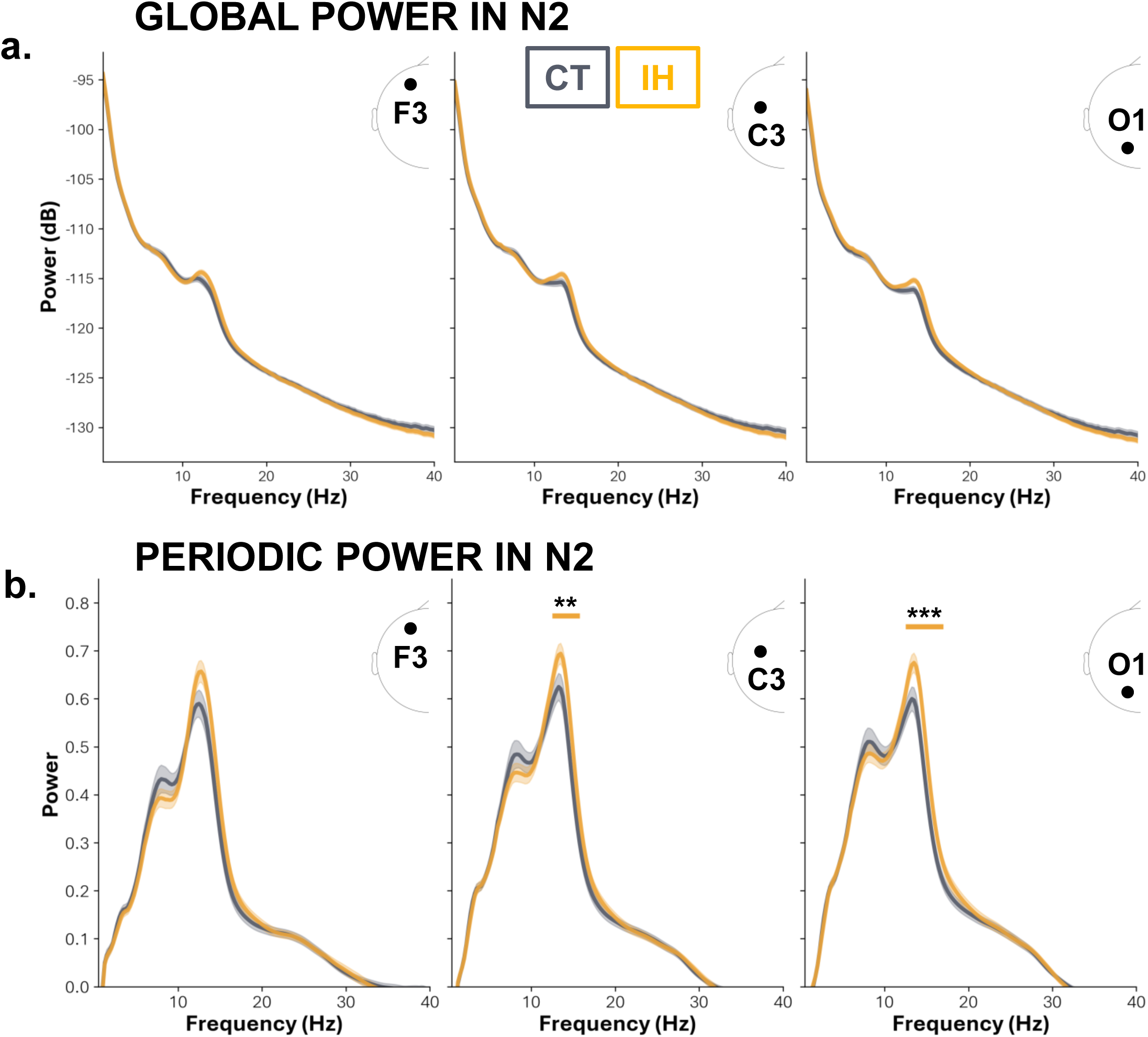
Participants with idiopathic hypersomnia have a higher power in the sigma band than controls. **a** Global, and **b** periodic power during N2 sleep, at channel F3, C3, and O1 (left, middle, and right respectively) in participants with idiopathic hypersomnia (yellow) compared to controls (grey). The yellow bar above the PSD in **b** at C3 and O1 indicates statistical differences. Statistics: linear mixed models with age and gender as covariate corrected with cluster-based multiple comparisons, p_CLUSTER_ < 0.05: *, p_CLUSTER_ < 0.01: **, p_CLUSTER_ < 0.001: ***. Shadings show the standard-error-of-the-mean (SEM) across participants.

### Spindle and Slow Wave Density

The increase in power in the sigma band was also reflected in the detection of sleep spindles: there was an increase in sleep spindle density in N2 sleep at the frontal (ß = 0.342, p = 0.008) and occipital electrodes (ß = 0.321, p = 0.008) in the IH group compared to control group (Figure 3a). There was an increase in slow waves in N3 sleep in the central (ß = 0.713, p = 0.047), and occipital electrode (ß = 1.165, p < 0.001) in the IH group compared to the control group (Figure 3b). The analysis of spindle clustering indicated that at F3 and O1, isolated spindles (cluster size = 1) were less frequent in the IH group than in the control group (Figure 4a, ß = -0.045, p < 0.001, and Supplemental Figure 4a, ß = -0.040, p < 0.001). At F3, pairs of spindles were more frequent in the IH group than in the control group (Figure 4a, ß = 0.026, p = 0.039). There was no group difference for clusters of sizes three, four, and above five spindles.

**Figure 3:**
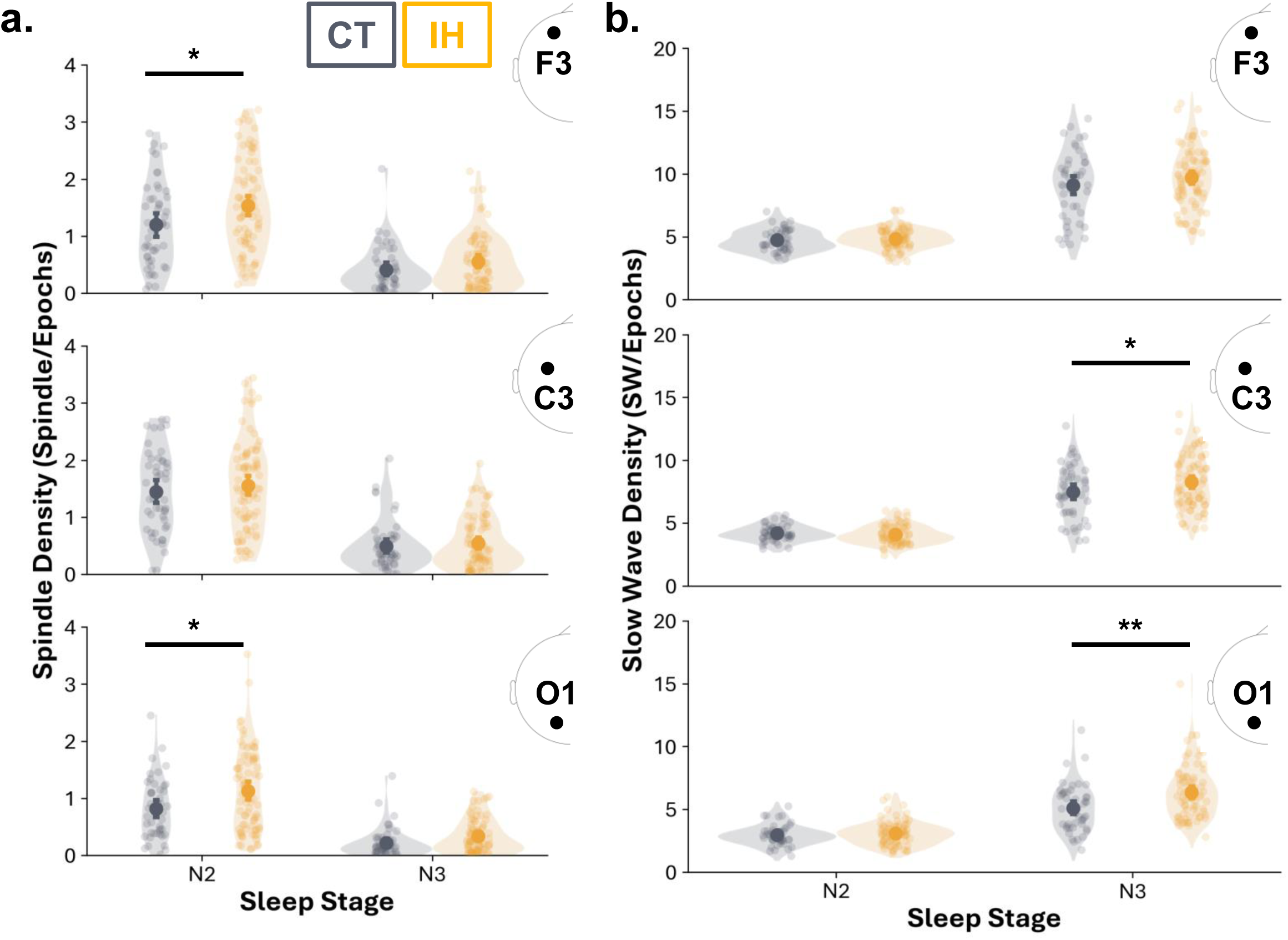
Sleep spindle and slow wave density were higher in IH in N2 and N3. **a** Sleep spindles density (number of sleep spindles divided by the number of epochs in a given sleep stage) in N2 and N3 for the IH (yellow) and control (grey) group. Small dots show individual data and large dots the population average. Error bars denote the standard error of the mean (SEM). **b** Slow wave density (number of slow waves divided by the number of epochs in a given sleep stage) in N2 and N3. F3 electrode on the top row, C3 in the middle, and O1 at the bottom. Statistics: linear mixed models with age and gender as covariate, with correction for multiple comparisons (false discovery rate), p < 0.05: *, p < 0.01: **, p < 0.001: ***.

**Figure 4:**
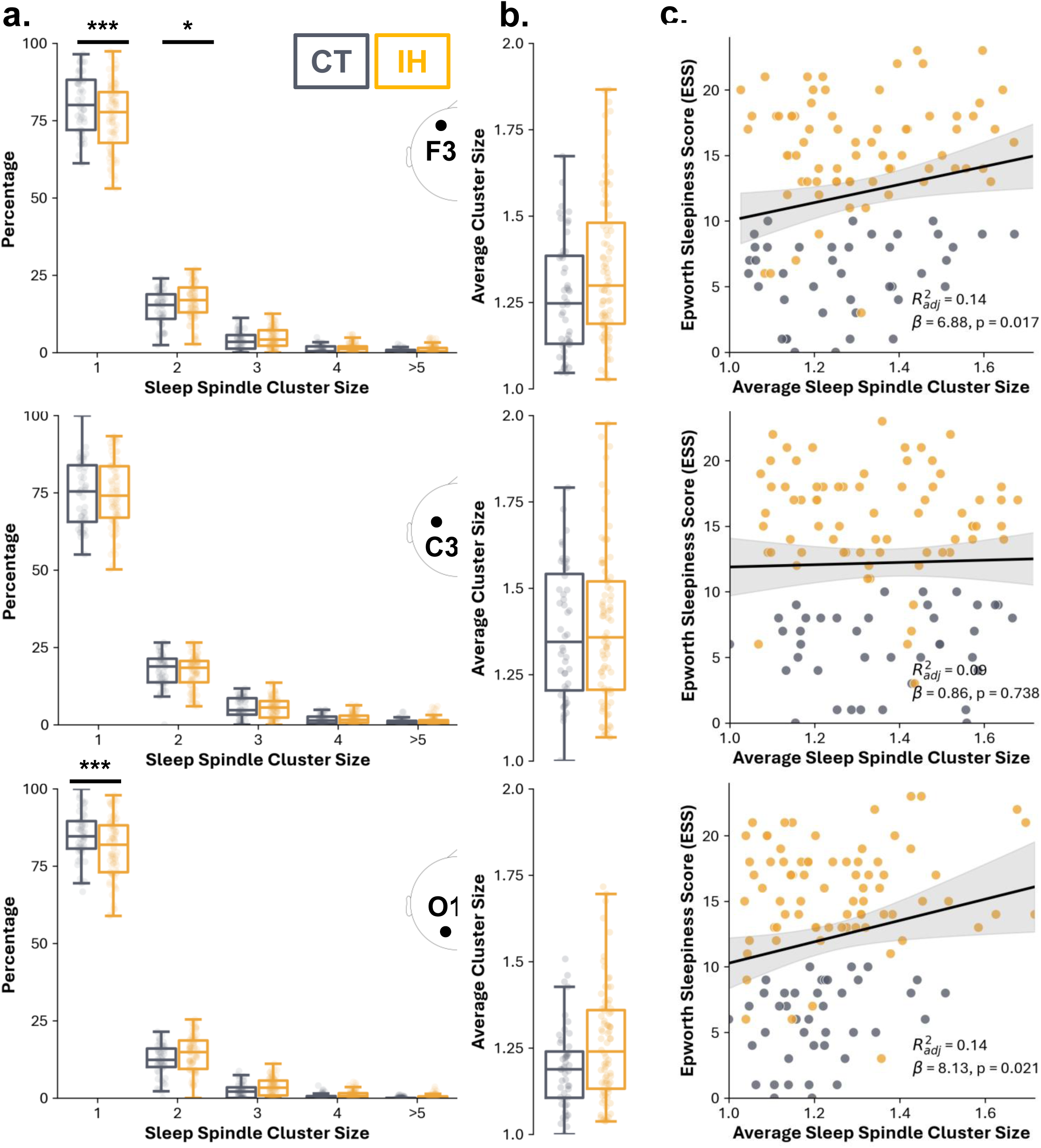
Sleep spindles were more clustered in the IH and predict the level of subjective sleepiness. **a** Percentage of sleep spindle clusters across size for the IH (yellow) and control (grey) groups. **b** Average sleep spindle cluster size per group. **c** Linear regression between average sleep spindle cluster size and Epworth sleepiness scale scores is represented by the solid black regression line (shading depicts the 95% CI). F3 electrode on the top row, C3 in the middle, and O1 at the bottom. Statistics for **a** and **b**: linear mixed models with age and gender as covariate with correction for multiple comparisons (false discovery rate), p < 0.05: *, p < 0.01: **, p < 0.001: ***. In **a** and **b**, box plots show the median (middle line), 1^st^ (bottom) and 3^rd^ (top) quartiles. Whiskers extend*s* to the most extreme non-outlier values (within 1.5×IQR). In all plots, dots show individual data points.

The results showed a non-significant higher average cluster size in IH group than in control group at F3 and O1 (Figure 4b, ß = 0.084, p = 0.55; ß = 0.071; p = 0.069 respectively). A linear regression analysis was conducted to examine the association between average spindle cluster size and Epworth sleepiness scale scores, controlling for age and gender. Higher average cluster sizes were associated with higher sleepiness scores at F3 and O1 (Figure 4c, ß = 6.88, SE = 2.84, t = 2.42, p = 0.011; ß = 8.13, SE = 3.48, t = 2.34, p = 0.011 respectively), with the model accounting for 12.5%, and 15% of the variance in sleepiness scores. The spindle cluster size in the IH participants with and without sleep drunkenness was similar (Supplemental Figure 4a, ß = 0.017, p = 0.720). We replicated previous findings showing an infraslow modulation (0.018 Hz) of the sigma power for F3, C3, and O1 electrodes (Figure 5). However, there was no differences between the IH group and the control group in any electrode (all p_CLUSTER_ > 0.05).

**Figure 5:**
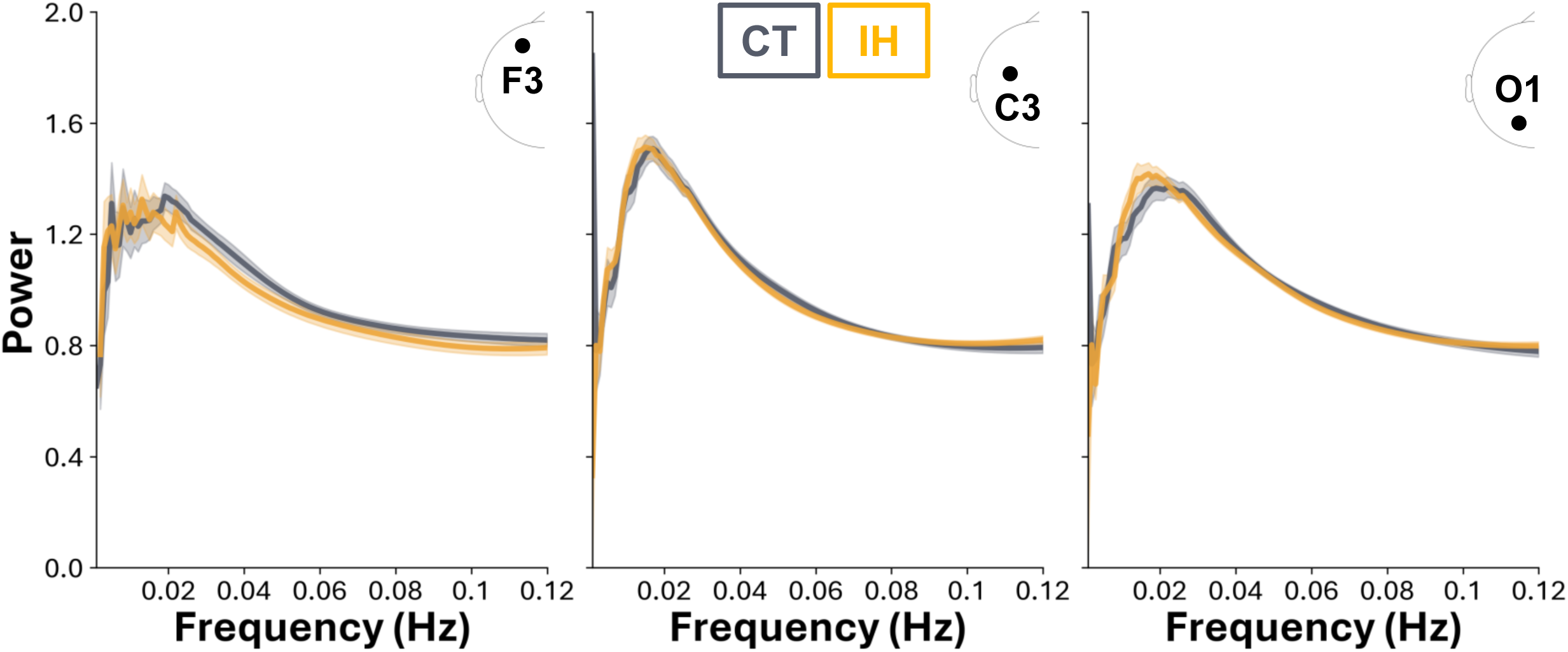
Infraslow sigma oscillations did not differ between groups. Normalized spectrum of 0.02 Hz oscillations in stable NREM sleep at F3 (left), C3 (center), and O1 (right) for the IH (yellow) and control (grey) groups. Shading area represents the standard error of the mean of the population per frequency bin. Statistics: linear mixed models with with age and gender as covariate cluster correction for multiple comparison, p_CLUSTER_ < 0.05: *, p_CLUSTER_ < 0.01: **, p_CLUSTER_ < 0.001: ***.

### Prediction of Idiopathic Hypersomnia Diagnosis via Sleep Metrics

Multivariate EEG -based models were developed to classify groups and predict a range of sleep quality metrics. As shown in Figure 6a (left panel), the area under the curve increased with the number of features, reaching a peak of approximately 0.81 for 11 features for the full sample and 0.74 for the balanced sample with 13 features (Supplementary Table 2). The red marker in Figure 6a indicates the optimal feature set (feature names in supplementary Table 2), while the accompanying confusion matrices (Figure 6a, middle and right panels) revealed that correct classification rates in the balanced dataset were significantly above chance-level (50%) but remained modest—69% for the control group and 64% for the IH group (Figure 6a, left panel). For the regression analyses, separate models were constructed to predict objective sleep measures (wake time after sleep onset and sleep efficiency), subjective sleep metrics (sleep duration during the weekend and score at the Epworth sleepiness scale), as well as total sleep time measured on night 2 and over 18 hours. In each case, features were sequentially added (up to 50 features), and model performance was evaluated using repeated 5-fold cross-validation (20 iterations). The optimal number of features was chosen based on the minimum RMSE, with error bars representing ± 1 standard deviation (Figure 6b–6d, left panels). Scatter plots comparing predicted versus true values (Figure 6b–6d, right panels) show a broad dispersion around the identity line, and the squared Spearman correlation coefficients (used as a proxy for R²) ranged from approximately 0.10 to 0.30 across the different metrics. These findings indicate that while the EEG-derived features are moderately associated with both objective and subjective sleep parameters, their overall predictive power remains limited. Collectively, these results demonstrate that the EEG-based modelling approach can capture some variability in sleep quality and subtype discrimination—yielding a good peak area under the curve (≈ 0.81; Figure 6a) and regression R² values between 0.10 and 0.30 (Figure.6b–6d)—but underscore the need for further model refinement and the incorporation of additional predictors to achieve clinically robust performance.

**Figure 6:**
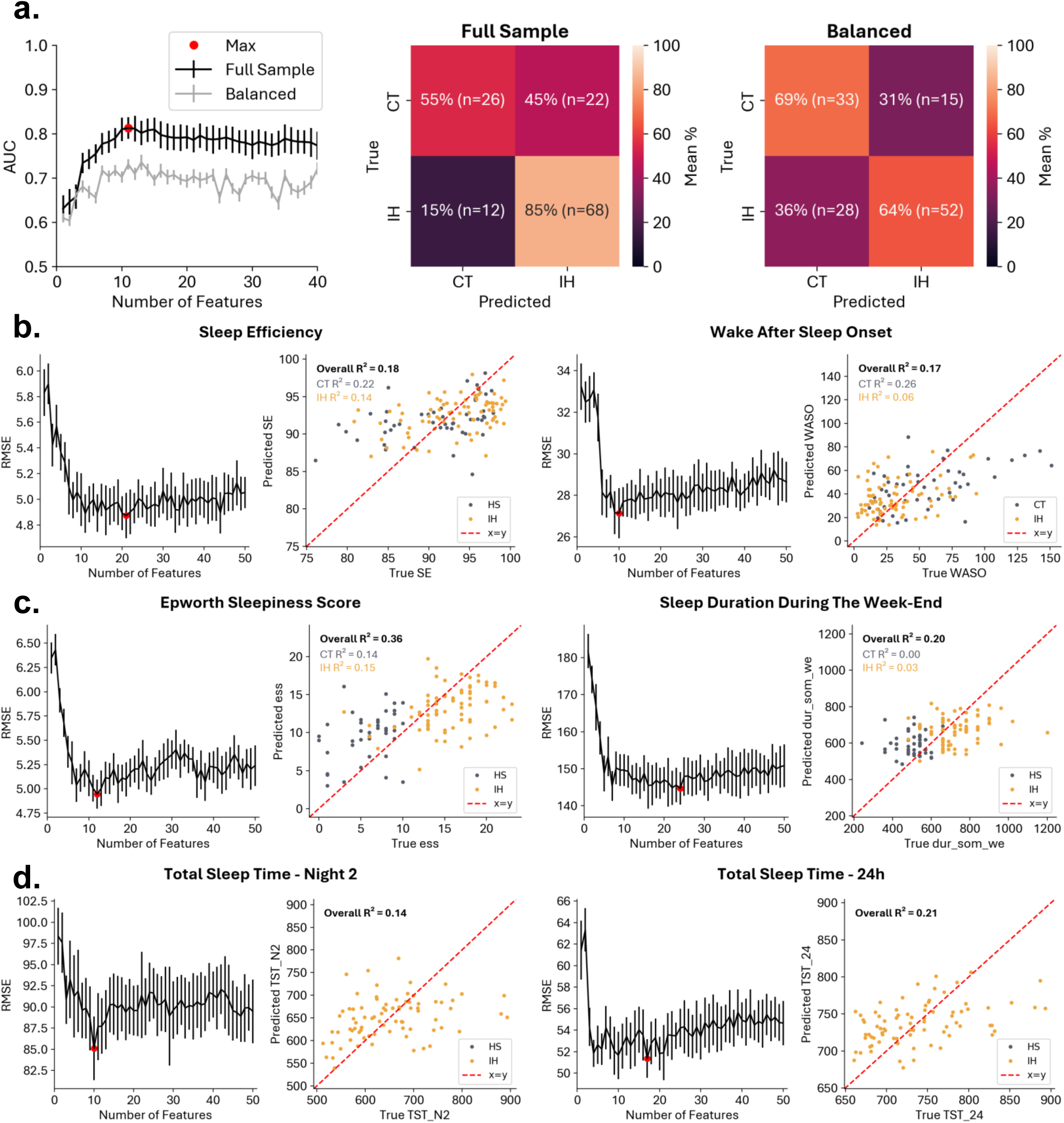
EEG-based features multivariate analyses predict group identify and sleep metrics. **a** Classifier results showing area under the curves and confusion matrices for both full (black) and balanced (dark grey) samples. The red marker highlights the optimal feature set, and confusion matrices display the percentage and count of correct classifications for control (grey) and idiopathic hypersomnia (yellow) groups. **b** Objective sleep measures (wake after sleep onset time and sleep efficiency) are evaluated via RMSE curves (left panels) and scatter plots (right panels) comparing predicted vs. true values. Data points are color-coded by group (control in grey, hypersomnia in yellow) with the red dashed line indicating the x=y identity line. **c** Subjective sleep metrics (sleep duration during the week and score at the Epworth sleepiness scale) are presented in the same format as (B). **d** Total sleep time predictions are shown for night 2 and the 24 hours (the latter for idiopathic hypersomnia only), again with RMSE curves and scatter plots. Error bars denote ±1 standard deviation across repeated cross-validation iterations.

## Discussion

In this controlled study of sleep EEG in 80 participants with IH with long sleep time, the percentage of N3 sleep and sleep efficiency were increased, while the percentage of REM sleep and wake after sleep onset were reduced in the IH group compared to the control group. Such an increase in the percentage of N3 sleep was also found in a previous study of 35 patients with IH^11^, although a meta-analysis of 10 previous studies (mixing IH participants with and without long sleep time) did not replicate this finding^10^. The restriction of our sample to IH with long sleep time, where well-consolidated sleep is expected (whereas sleep in IH with normal sleep time may be disrupted, as in narcolepsy type 2), may explain this slight discrepancy. However, the high percentage of N3 sleep compared to the low percentage of REM sleep in our IH group may also reflect the way the sleep test was performed in our sleep clinic. Participants were awoken at 6:30 am before the first MSLT. This procedure could have shortened the total sleep time in IH (normalizing its duration compared to controls) and reduced the amount of REM sleep (which increases at the end of the night) in favor of N3 sleep.

Beyond the absolute or relative quantity of sleep stages, we examined qualitative differences across groups by extracting and analyzing hypnodensities. Group-level hypnodensity profiles were broadly comparable between IH and control groups, except for more mixed wake/N1 sleep epochs in IH. This finding contrasts with similar analyses in narcolepsy type 1, where dominant probabilities were more mixed with other sleep stages, suggesting more hybrid stages^19^. Still, we found that wakefulness after sleep onset (intra-sleep wakefulness) had a higher probability of N1 in IH individuals. These intrusions of sleep into wakefulness may be a marker of the difficulty of fully awakening from sleep in IH individuals. Analysis of epoch-to-epoch dynamics showed that consecutive NREM sleep epochs were less similar in the IH than in control recordings, suggesting greater moment-to-moment variability. Since there was no increase in hybrid sleep stages and since hypnograms did not reveal more awakenings in IH, this variability does not seem to result from a greater instability as recently shown in insomnia^21^. To further explore the dynamics of sleep in IH, it would be interesting to consider other markers of sleep instability such as CAP and compare the rate of CAP between IH and controls^13^.

In terms of microstructural analysis, increased sleep spindles and slow wave densities in IH were two notable findings. Here, IH participants had increased sigma power and a higher density of sleep spindles. Sleep spindles were also more clustered in IH compared to controls. These results confirm and extend the previous findings of two research groups, who found a higher density of sleep spindles during N2 sleep in IH than in controls and narcolepsy^16,17^. This increased activity and clustering of spindle, may reflect an over-stabilization of thalamocortical networks, which may underlie the pathophysiology of IH. As previously suggested, this mechanism may contribute to severe sleep inertia^16,17^. Moreover, given that periods with a higher density of sleep spindles have been associated with reduced sensitivity to external noise during sleep^44^, an excess or abnormal clustering of spindles in IH might further diminish cortical responsiveness and impair efficient arousal mechanisms, potentially explaining sleep drunkenness. Importantly, these changes in spindle activity were not reflected in the infraslow rhythmic modulation of sigma power, suggesting that long-timescale regulatory mechanisms of sleep under the influence of noradrenergic activity could be similar in IH and controls^24^.

Regarding delta power in IH, previous studies have yielded inconsistent results, with an increase over night in some studies (including participants with IH and some psychiatric disorders) on one hand^10,45^, and a decrease in a study of 10 patients with IH without long sleep time on the other hand^14^. In our IH series, while a global spectral analysis did not reveal differences in overall delta power compared to controls, a targeted analysis of slow waves patterns during N3 sleep revealed higher slow waves density than in the control group. This finding suggests that global delta power measures may obscure specific microarchitectural changes in slow wave activity. The fact that delta power does not just reflect the number but also amplitude of slow waves could explain differences between slow waves density and delta power. Importantly, an increase in slow wave density in IH patients could promote sleep drunkenness in these patients, as the prevalence of slow waves is associated with sleep inertia^46^. One potential mechanism underlying this increase could be an enhanced homeostatic sleep regulation^47^. An elevated sleep drive may stem from a greater restorative need, manifesting as an increased slow wave activity. Indeed, the synaptic homeostasis hypothesis proposes that slow waves activity plays a crucial role in downscaling synaptic strength accumulated during wakefulness, with higher slow waves density reflecting an amplified need for synaptic renormalization^48^. It is possible that individuals with IH tend to create and strengthen synaptic contact at a higher rate during wakefulness, which would lead to an increase in slow waves during sleep as proposed for other sleep disorders^49^. Alternatively, changes in brain metabolism have also been associated with an increase in slow waves and could represent the cause or source of the enhanced slow wave activity observed here during sleep^50^. Both hypotheses suggest that the ultimate cause of IH symptoms could be found during wakefulness rather than sleep, in line with our findings showing a mostly preserved sleep in IH.

The classification of IH is based on the exclusion of other hypersomnolence disorders, such as narcolepsy type 1, insufficient sleep syndrome, or other sleep disorders. Accordingly, the clinical categorization of IH based on conventional sleep measures remains inconsistent when compared to normal sleepers, with significant variability between studies^11,14,14,51–55^. This may be because IH is characterized by many small differences that are difficult to detect in isolation and in small samples. In this context, our findings show that, when integrating many indexes derived from the first night PSG recordings (but not MSLT latency or total sleep time during 24h recordings), our EEG-based model achieved a good balanced classification performance with an AUC of 0.74 and a correct classification rate around 65%. Our approach could therefore represent an interesting aid for the diagnosis of IH, based on a single night of PSG recordings. Yet, the predictive power of our models for objective sleep measures remained low, suggesting that a significant proportion of the inter-individual variability remains unaccounted for. Our results thus support the notion that leveraging a combination of EEG features rather than relying on conventional sleep staging or a small set of macrostructural sleep parameters may be key to refining our understanding of sleep disorders^18,56,57^.

Our study has several limitations, including EEG recordings limited to 3 channels (which limits the spatial analysis of EEG), and a control group smaller than the IH group. On the other hand, the IH group analyzed here was large (although IH is a rare disorder), well-characterized (using not only MLST but also long-term bed rest recordings), untreated, without any individuals with possible psychiatric hypersomnia, resulting in homogeneity and robustness in the results. In addition, measurements were adjusted for sex and age.

In conclusion, our findings show that IH with long sleep time is characterized by distinct but subtle EEG signatures that paradoxically suggest a seemingly ‘better-than-normal’ sleep structure. This dissociation between normal nocturnal sleep microarchitecture and daytime hypersomnolence underscores the need to further explore daytime neural mechanisms and suggests a potentially deficient arousal system in IH. Ultimately, these findings may lead to a refinement of the diagnostic criteria for IH.

## Acknowledgements

We thank the medical and para-medical staff for their never-stopping care for the patients, their scrutinous work for the acquisition and scoring of sleep recordings, and the sharing of their clinical expertise. ALC was supported by a grant from Ecole Doctorale Frontières de l’Innovation en Recherche et Education–Programme Bettencourt. TA was supported by a grant from the Agence Nationale pour la Recherche (ANR-22-CE37–0006-01) and from the European Research Council (ERC-StG SleepingAwake 101116748). DO was supported by a grant from the European Research Council (ERC-Cog Creadoze 101087031).

## Glossary

CAP: cyclic alternating pattern
FFT: fast Fourier transformation
IH: idiopathic hypersomnia
MSLT: multiple sleep latency test
PSD: Power Spectral Density

## References

1. Sateia MJ. International Classification of Sleep Disorders-Third Edition. Chest. 2014;146(5):1387–1394. 10.1378/chest.14-0970

2. Šonka K, Šusta M, Billiard M. Narcolepsy with and without cataplexy, idiopathic hypersomnia with and without long sleep time: a cluster analysis. Sleep Med. 2015;16(2):225–231. doi:10.1016/j.sleep.2014.09.016

3. Billiard M, Sonka K. Idiopathic hypersomnia. Sleep Medicine Reviews. 2016;29:23-33. doi:10.1016/j.smrv.2015.08.007

4. Fronczek R, Arnulf I, Baumann CR, Maski K, Pizza F, Trotti LM. To split or to lump? Classifying the central disorders of hypersomnolence. Sleep. 2020;43(8):zsaa044. doi:10.1093/sleep/zsaa044

5. Gool JK, Zhang Z, Oei MSSL, et al. Data-Driven Phenotyping of Central Disorders of Hypersomnolence With Unsupervised Clustering. Neurology. 2022;98(23):e2387–e2400. doi:10.1212/WNL.0000000000200519

6. Plante DT, Hagen EW, Barnet JH, Mignot E, Peppard PE. Prevalence and Course of Idiopathic Hypersomnia in the Wisconsin Sleep Cohort Study. Neurology. 2024;102(2):e207994. doi:10.1212/WNL.0000000000207994

7. Lippert J, Halfter H, Heidbreder A, et al. Altered dynamics in the circadian oscillation of clock genes in dermal fibroblasts of patients suffering from idiopathic hypersomnia. PLoS One. 2014;9(1):e85255. doi:10.1371/journal.pone.0085255

8. Materna L, Halfter H, Heidbreder A, et al. Idiopathic Hypersomnia Patients Revealed Longer Circadian Period Length in Peripheral Skin Fibroblasts. Front Neurol. 2018;9:424. doi:10.3389/fneur.2018.00424

9. Rye DB, Bliwise DL, Parker K, et al. Modulation of vigilance in the primary hypersomnias by endogenous enhancement of GABAA receptors. Sci Transl Med. 2012;4(161):161ra151. doi:10.1126/scitranslmed.3004685

10. Plante DT, Cook JD, Barbosa LS, et al. Establishing the objective sleep phenotype in hypersomnolence disorder with and without comorbid major depression. Sleep. 2019;42(6):zsz060. doi:10.1093/sleep/zsz060

11. Evangelista E, Lopez R, Barateau L, et al. Alternative diagnostic criteria for idiopathic hypersomnia: A 32-hour protocol. Ann Neurol. 2018;83(2):235–247. doi:10.1002/ana.25141

12. Vernet C, Arnulf I. Idiopathic Hypersomnia with and without Long Sleep Time: A Controlled Series of 75 Patients. Sleep. 2009;32(6):753–759. doi:10.1093/sleep/32.6.753

13. Pizza F, Ferri R, Poli F, Vandi S, Cosentino FII, Plazzi G. Polysomnographic study of nocturnal sleep in idiopathic hypersomnia without long sleep time. J Sleep Res. 2013;22(2):185–196. doi:10.1111/j.1365-2869.2012.01061.x

14. Sforza E, Gaudreau H, Petit D, Montplaisir J. Homeostatic sleep regulation in patients with idiopathic hypersomnia. Clinical Neurophysiology. 2000;111(2):277–282. doi:10.1016/S1388-2457(99)00242-4

15. Cairns A, Bogan R. Comparison of the macro and microstructure of sleep in a sample of sleep clinic hypersomnia cases. Neurobiology of Sleep and Circadian Rhythms. 2019;6:62–69. doi:10.1016/j.nbscr.2019.02.001

16. Bové A, Culebras A, Moore JT, Westlake RE. Relationship between sleep spindles and hypersomnia. Sleep. 1994;17(5):449–455. doi:10.1093/sleep/17.5.449

17. Delrosso LM, Chesson AL, Hoque R. Manual characterization of sleep spindle index in patients with narcolepsy and idiopathic hypersomnia. Sleep Disord. 2014;2014:271802. doi:10.1155/2014/271802

18. Stephan AM, Siclari F. Reconsidering sleep perception in insomnia: from misperception to mismeasurement. J Sleep Res. 2023;32(6):e14028. doi:10.1111/jsr.14028

19. Stephansen JB, Olesen AN, Olsen M, et al. Neural network analysis of sleep stages enables efficient diagnosis of narcolepsy. Nat Commun. 2018;9(1):5229. doi:10.1038/s41467-018-07229-3

20. Vallat R, Walker MP. An open-source, high-performance tool for automated sleep staging. Elife. 2021;10:e70092. doi:10.7554/eLife.70092

21. Herzog R, Crosbie F, Aloulou A, et al. A continuous approach to explain insomnia and subjective-objective sleep discrepancy. Commun Biol. 2025;8(1):1–14. doi:10.1038/s42003-025-07794-6

22. Donoghue T, Haller M, Peterson EJ, et al. Parameterizing neural power spectra into periodic and aperiodic components. Nat Neurosci. 2020;23(12):1655–1665. doi:10.1038/s41593-020-00744-x

23. Osorio-Forero A, Cardis R, Vantomme G, et al. Noradrenergic circuit control of non-REM sleep substates. Current Biology. 2021;31(22):5009–5023.e7. doi:10.1016/j.cub.2021.09.041

24. Osorio-Forero A, Foustoukos G, Cardis R, et al. Infraslow noradrenergic locus coeruleus activity fluctuations are gatekeepers of the NREM–REM sleep cycle. Nat Neurosci. 2025;28(1):84–96. doi:10.1038/s41593-024-01822-0

25. Foustoukos G, Lüthi A. Monoaminergic signaling during mammalian NREM sleep - Recent insights and next-level questions. Current Opinion in Neurobiology. 2025;92:103025. doi:10.1016/j.conb.2025.103025

26. Arnulf I, Leu-Semenescu S, Dodet P. Precision Medicine for Idiopathic Hypersomnia. Sleep Med Clin. 2022;17(3):379–398. doi:10.1016/j.jsmc.2022.06.016

27. Troester MM, Quan SF, Medicine AA of S, Berry RB. The AASM Manual for the Scoring of Sleep and Associated Events, Version 3. American Academy Of Sleep Medicine; 2023. https://books.google.fr/books?id=uKSczwEACAAJ

28. Berry RB, Brooks R, Gamaldo C, et al. AASM Scoring Manual Updates for 2017 (Version 2.4). Journal of Clinical Sleep Medicine. 2017;13(05):665–666. doi:10.5664/jcsm.6576

29. Nikkonen S, Somaskandhan P, Korkalainen H, et al. Multicentre sleep-stage scoring agreement in the Sleep Revolution project. Journal of Sleep Research. 2024;33(1):e13956. doi:10.1111/jsr.13956

30. Norman RG, Pal I, Stewart C, Walsleben JA, Rapoport DM. Interobserver Agreement Among Sleep Scorers From Different Centers in a Large Dataset. Sleep. 2000;23(7):1-8. doi:10.1093/sleep/23.7.1e

31. Hanif U, Aloulou A, Crosbie F, et al. Deciphering Insomnia: Benchmarking Automated Sleep Staging Algorithms for Complex Sleep Disorders. Journal of Sleep Research. n/a(n/a):e70048. doi:10.1111/jsr.70048

32. Pedregosa F, Varoquaux G, Gramfort A, et al. Scikit-learn: Machine Learning in Python. MACHINE LEARNING IN PYTHON. Published online October 2011.

33. Cover TM, Thomas JA. ELEMENTS OF INFORMATION THEORY. Published online 2005.

34. Huijben IAM, Hermans LWA, Rossi AC, Overeem S, van Gilst MM, van Sloun RJG. Interpretation and further development of the hypnodensity representation of sleep structure. Physiol Meas. 2023;44(1):015002. doi:10.1088/1361-6579/aca641

35. Gramfort A, Luessi M, Larson E, et al. MEG and EEG data analysis with MNE-Python. Front Neurosci. 2013;7:267. doi:10.3389/fnins.2013.00267

36. Virtanen P, Gommers R, Oliphant TE, et al. SciPy 1.0: fundamental algorithms for scientific computing in Python. Nat Methods. 2020;17(3):261–272. doi:10.1038/s41592-019-0686-2

37. Champetier P, André C, Weber FD, et al. Age-related changes in fast spindle clustering during non- rapid eye movement sleep and their relevance for memory consolidation. Sleep. 2023;46(5):zsac282. doi:10.1093/sleep/zsac282

38. Lecci S, Fernandez LMJ, Weber FD, et al. Coordinated infraslow neural and cardiac oscillations mark fragility and offline periods in mammalian sleep. Science Advances. 2017;3(2):e1602026. doi:10.1126/sciadv.1602026

39. Ding C, Peng H. Minimum redundancy feature selection from microarray gene expression data. In: Computational Systems Bioinformatics. CSB2003. Proceedings of the 2003 IEEE Bioinformatics Conference. CSB2003.; 2003:523-528. doi:10.1109/CSB.2003.1227396

40. Chen T, Guestrin C. XGBoost: A Scalable Tree Boosting System. In: Proceedings of the 22nd ACM SIGKDD International Conference on Knowledge Discovery and Data Mining. ACM; 2016:785-794. doi:10.1145/2939672.2939785

41. Seabold S, Perktold J. Statsmodels: Econometric and Statistical Modeling with Python. *scipy*. Published online May 1, 2010. doi:10.25080/Majora-92bf1922-011

42. Benjamini Y, Hochberg Y. Controlling the False Discovery Rate: A Practical and Powerful Approach to Multiple Testing. Journal of the Royal Statistical Society: Series B (Methodological). 1995;57(1):289–300. doi:10.1111/j.2517-6161.1995.tb02031.x

43. Maris E, Oostenveld R. Nonparametric statistical testing of EEG- and MEG-data. J Neurosci Methods. 2007;164(1):177–190. doi:10.1016/j.jneumeth.2007.03.024

44. Dang-Vu TT, McKinney SM, Buxton OM, Solet JM, Ellenbogen JM. Spontaneous brain rhythms predict sleep stability in the face of noise. Curr Biol. 2010;20(15):R626–627. doi:10.1016/j.cub.2010.06.032

45. Walacik-Ufnal E, Piotrowska AJ, Wołyńczyk-Gmaj D, et al. Narcolepsy type 1 and hypersomnia associated with a psychiatric disorder show different slow wave activity dynamics. Acta Neurobiol Exp (Wars). 2017;77(2):147–156. doi:10.21307/ane-2017-047

46. Trotti LM. Waking up is the hardest thing I do all day: Sleep inertia and sleep drunkenness. Sleep Med Rev. 2017;35:76–84. doi:10.1016/j.smrv.2016.08.005

47. Tononi G, Cirelli C. Sleep function and synaptic homeostasis. Sleep Med Rev. 2006;10(1):49-62. doi:10.1016/j.smrv.2005.05.002

48. Tononi G, Cirelli C. Sleep and the price of plasticity: from synaptic and cellular homeostasis to memory consolidation and integration. Neuron. 2014;81(1):12–34. doi:10.1016/j.neuron.2013.12.025

49. Avvenuti G, Bernardi G. Chapter 3 - Local sleep: A new concept in brain plasticity. In: Quartarone A, Ghilardi MF, Boller F, eds. Handbook of Clinical Neurology. Vol 184. Neuroplasticity. Elsevier; 2022:35-52. doi:10.1016/B978-0-12-819410-2.00003-5

50. Krueger JM, Nguyen JT, Dykstra-Aiello CJ, Taishi P. Local sleep. Sleep Medicine Reviews. 2019;43:14-21. doi:10.1016/j.smrv.2018.10.001

51. Maski KP, Colclasure A, Little E, et al. Stability of nocturnal wake and sleep stages defines central nervous system disorders of hypersomnolence. Sleep. 2021;44(7):zsab021. doi:10.1093/sleep/zsab021

52. Plante DT. Nocturnal sleep architecture in idiopathic hypersomnia: a systematic review and meta- analysis. Sleep Medicine. 2018;45:17–24. doi:10.1016/j.sleep.2017.10.005

53. Sforza E, Roche F, Barthélémy JC, Pichot V. Diurnal and nocturnal cardiovascular variability and heart rate arousal response in idiopathic hypersomnia. Sleep Medicine. 2016;24:131–136. doi:10.1016/j.sleep.2016.07.012

54. Vanková J, Nevsímalová S, Sonka K, Spacková N, Svejdová-Blazejová K. Increased REM density in narcolepsy-cataplexy and the polysymptomatic form of idiopathic hypersomnia. Sleep. 2001;24(6):707–711. doi:10.1093/sleep/24.6.707

55. Vgontzas AN, Bixler EO, Kales A, Criley C, Vela-Bueno A. Differences in nocturnal and daytime sleep between primary and psychiatric hypersomnia: diagnostic and treatment implications. Psychosom Med. 2000;62(2):220–226. doi:10.1097/00006842-200003000-00013

56. Andrillon T, Solelhac G, Bouchequet P, et al. Revisiting the value of polysomnographic data in insomnia: more than meets the eye. Sleep Med. 2020;66:184–200. doi:10.1016/j.sleep.2019.12.002

57. Herzog R, Crosbie F, Aloulou A, et al. Sleep and wake intrusions: A continuous approach to explain insomnia and sleep state misperception. Published online October 18, 2024. doi:10.21203/rs.3.rs-4924650/v1

