## Supplemental Figure 1 for "EEG Signature of Idiopathic Hypersomnia: Insights from Sleep Microarchitecture and Hypnodensity Metrics"

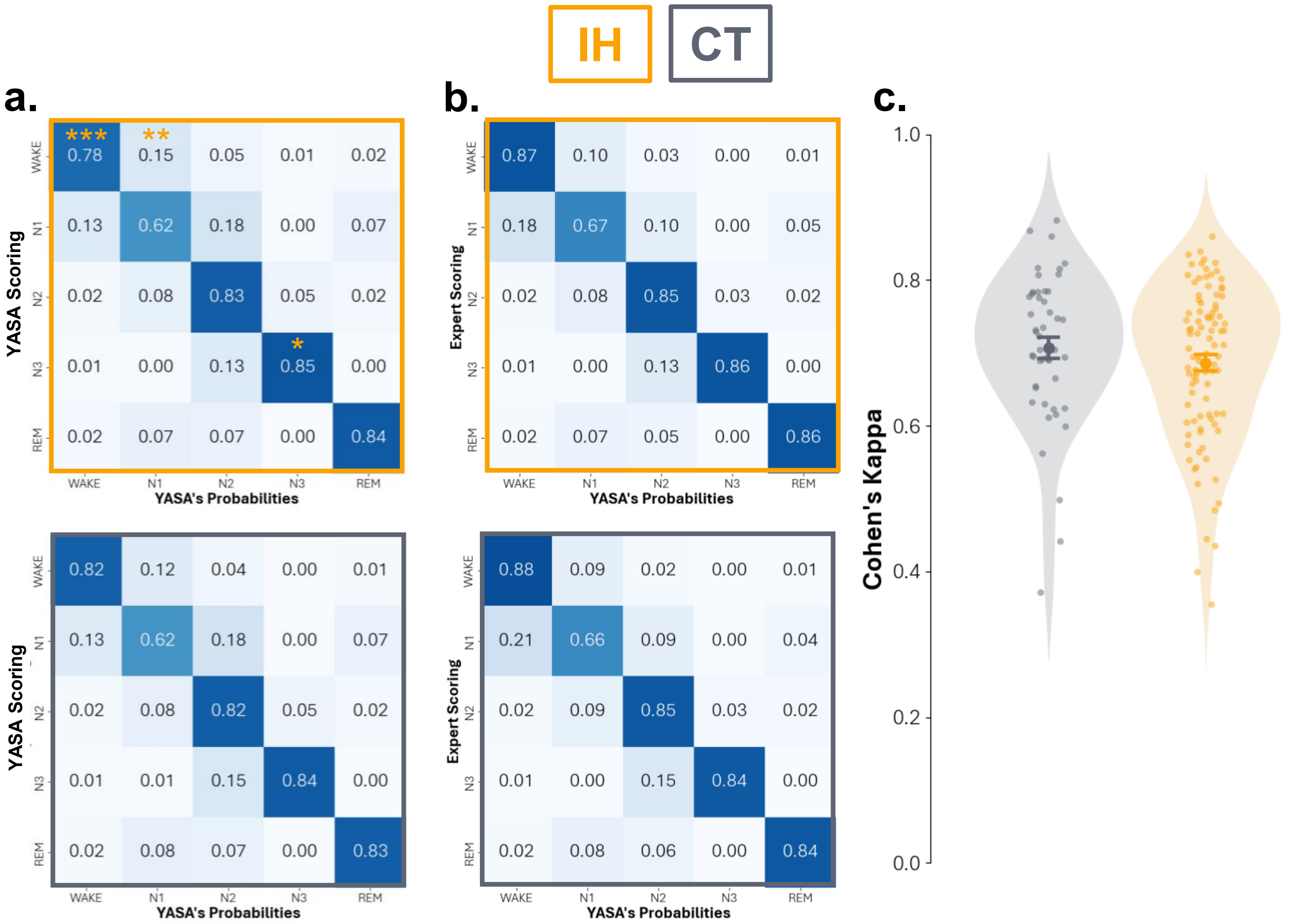


**Supplemental Figure 1: Hypnodensity matrices for the automated and human scoring.** Hypnodensities according to the sleep stage scored by **a** YASA, and **b** experts of participants with IH on the upper part (yellow squares) and controls on the lower part (grey squares). **c** Cohen's Kappa of YASA automatic sleep staging against expert scorers. Participants with IH in yellow compared to controls in grey. Statistics: Linear Mixed Models with age and gender as covariate correction for multiple comparison (false discovery rate), $p < 0.05$: *, $p < 0.01$: **, $p < 0.001$: ***.
