## Supplemental Figure 2 for "EEG Signature of Idiopathic Hypersomnia: Insights from Sleep Microarchitecture and Hypnodensity Metrics"

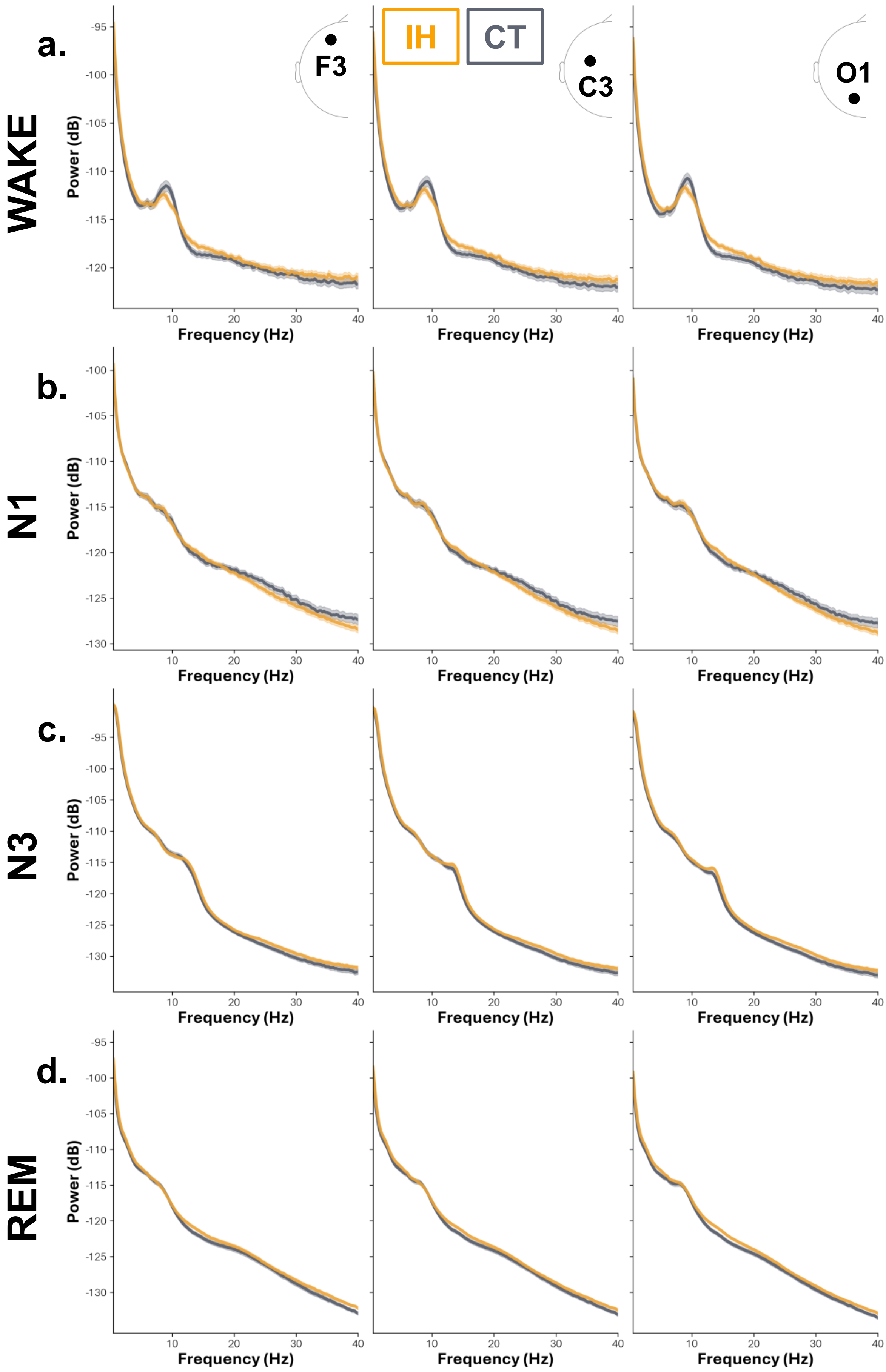


**Supplemental Figure 2: Global power in wake during the night, N1, N3 and REM sleep.** Global power in **a** wake during the night, **b** N1, **c** N3, and **d** REM sleep at channel F3, C3, and O1 (left, middle, and right, respectively) in participants with IH (yellow) compared to controls (grey). Shading area represents the standard error of the mean of the population per frequency bin.
