## Supplemental Figure 3 for "EEG Signature of Idiopathic Hypersomnia: Insights from Sleep Microarchitecture and Hypnodensity Metrics"

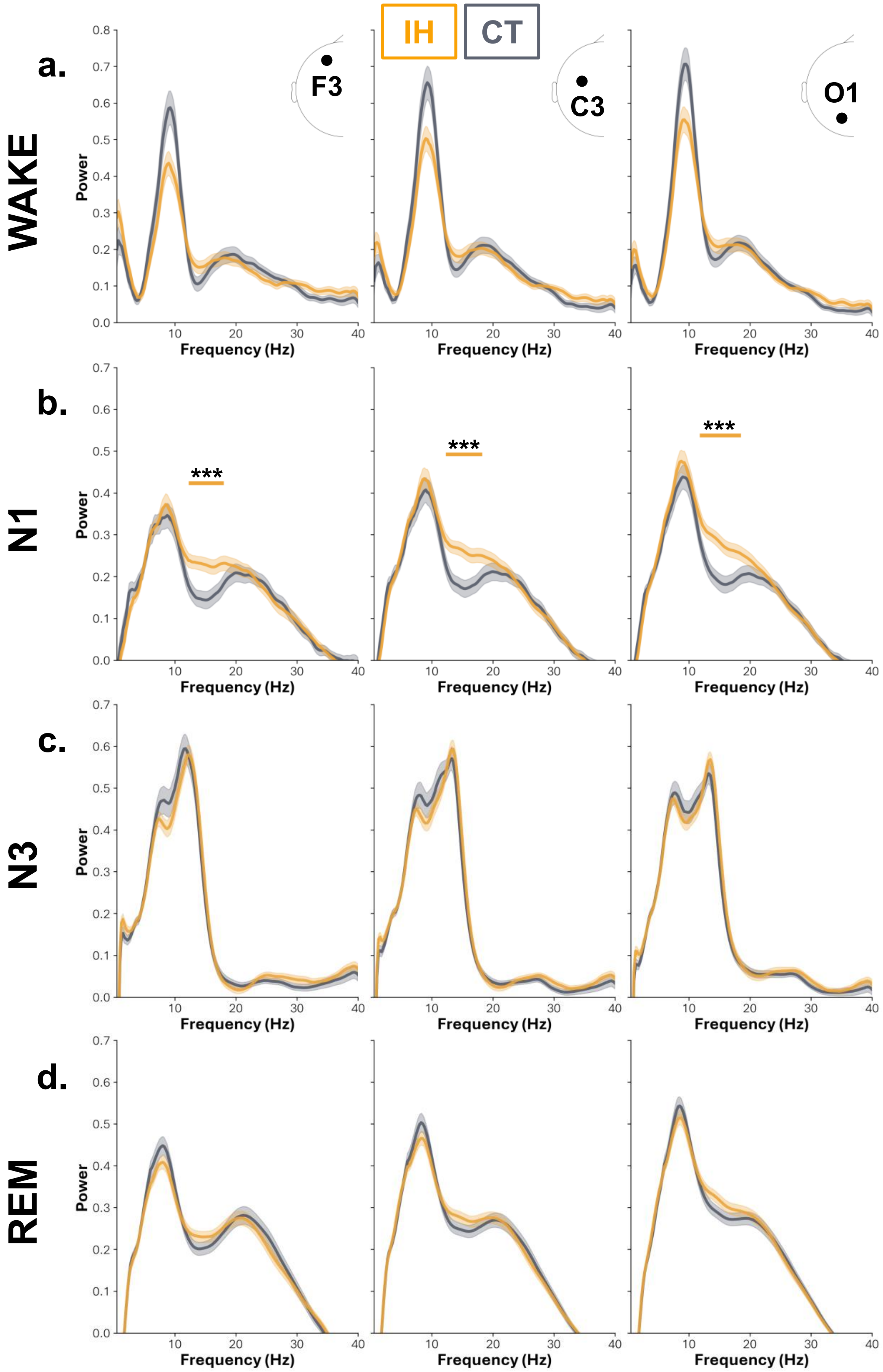


**Supplemental Figure 3: Periodic power in wake during the night, N1, N3 and REM sleep.** Periodic power in **a** wake during the night, **b** N1, **c** N3, and **d** REM sleep at channel F3, C3, and O1 (left, middle, and right, respectively) in participants with IH (yellow) compared to controls (grey). The yellow bar above the PSD in **b** indicates statistical differences. Shading area represents the standard error of the mean of the population per frequency bin. Statistics: linear mixed models with age and gender as covariate corrected with cluster correction for multiple comparisons, $p_{CLUSTER} < 0.05$: *, $p_{CLUSTER} < 0.01$: **, $p_{CLUSTER} < 0.001$: ***.
