## Supplemental Figure 4 for "EEG Signature of Idiopathic Hypersomnia: Insights from Sleep Microarchitecture and Hypnodensity Metrics"

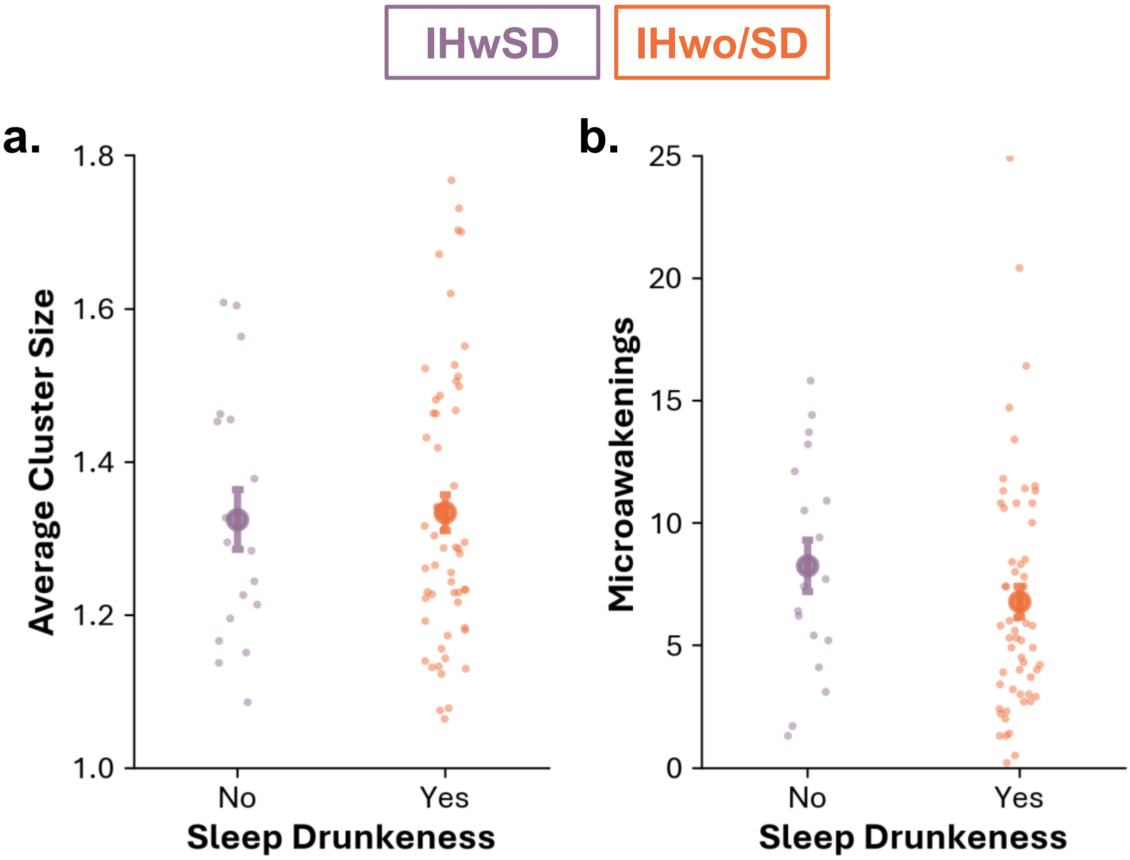


**Supplemental Figure 4: Spindle cluster size and microawakenings for IH with (IHwSD) and without sleep drunkenness (IHwo/SD). a** Average cluster size and **b** microawakening index in IH with (purple) and without (orange) sleep drunkenness. Statistics: linear mixed models with age and gender as covariate, corrected for multiple comparisons (false rate discovery), $p < 0.05$: *, $p < 0.01$: **, $p < 0.001$: ***.
