## Supplemental Table 1 for "EEG Signature of Idiopathic Hypersomnia: Insights from Sleep Microarchitecture and Hypnodensity Metrics"

*Supplemental Table 1: Demographic, clinical and sleep measures in participants with idiopathic hypersomnia with long sleep time and controls.*

| Category | Feature | Definition / Unit | Channels | Sleep stage(s) |
|---|---|---|---|---|
| **Event counts** | Slow wave count | Number of events detected | F3, C3, O1 | All (Wake, N1, N2, N3, REM) |
| | Sleep spindle count | | | N2 and N3 |
| **Event density** | Slow wave density | Events per 30-s epochs | F3, C3, O1 | All |
| | Sleep spindle density | | | N2 and N3 |
| **Amplitude / morphology** | Slow wave and sleep spindle amplitude | Peak-to-peak (µV) | F3, C3, O1 | All |
| | Slow wave downward and upward slope | µV·s-1 | | All |
| | Sleep spindle duration | duration (s) | | N2 and N3 |
| | Sleep spindle frequency | Median instantaneous frequency of spindle (in Hz) | | N2 and N3 |
| | Slow wave frequency | Hz | | All |
| **Spectral Absolute power** | Absolute power in the delta, theta, alpha, sigma and beta frequency ranges | $\mu V^2/Hz$ | F3, C3, O1 | All |
| | Sleep spindle absolute power | Median absolute power (in log10 $\mu V^2$) | | |
| **Spectral Relative Power** | Relative power in the delta, theta, alpha, sigma and beta frequency ranges | % of total power 0.5 to 40Hz | F3, C3, O1 | All |
| | Sleep Spindle Relative Power | Median % [11.25; 15.75Hz] of total power 1 to 30 | | N2 and N3 |
| **Aperiodic Fit** | Exponent, Offset | 1/F Slope and intercept | F3, C3, O1 | All |
| **Hypnodensity-derived features** | dKL, Entropy | | C3, Chin EMG, Left EOG | All |
