## Supplemental Table 2 for "EEG Signature of Idiopathic Hypersomnia: Insights from Sleep Microarchitecture and Hypnodensity Metrics"

*Supplemental Table 2: Top 13 mRMR-Ranked EEG Features for Classifying IH and HS subjects.*

| Rank | EEG Feature name | Stage | Electrode |
|---|---|---|---|
| 1 | dKL | N3 | |
| 2 | Sleep Spindle Count | N3 | O1 |
| 3 | Slow Wave Count | REM | C3 |
| 4 | Slow Wave Density | N3 | O1 |
| 5 | Absolute Beta Power | N3 | O1 |
| 6 | Hypnodensities Entropy | N1 | |
| 7 | Sleep Spindle Frequency | N2 | F3 |
| 8 | dKL | N2 | |
| 9 | Sleep Spindle Frequency | N3 | F3 |
| 10 | Slow Wave Count | N3 | O1 |
| 11 | Slow Wave Frequency | N1 | O1 |
| 12 | Absolute Sigma Power | REM | C3 |
| 13 | Sleep Spindle Count | N2 | O1 |
